# A novel purity analysis method of bovine milk-derived exosomes by two-dimensional high-performance liquid chromatography

**DOI:** 10.1101/2023.02.22.529450

**Authors:** Lu Lu, Chunle Han, Quan Zhang, Miao Wang, Dongli Qi, Mengya Gao, Na Wang, Jianxin Yin, Fengwei Dong, Xiaohu Ge

## Abstract

Exosomes have been implicated in many biological processes as intercellular communication carriers. Because exosomes are increasingly explored as natural vehicles for cell and gene therapies, and drug delivery applications, it is critical to have high-quality samples. Protein:particle ratio, ELISA, western blotting, polymerase chain reaction, and size-exclusion high-performance liquid chromatography are the main methods that have been extensively applied to analyze exosomes purity in recent years. However, there is currently no well-established method that is convenient for routine quality analysis of small-size impurities in exosomes samples. Here, a size-exclusion high-performance liquid chromatography (SE-HPLC), an ion-exchange high-performance liquid chromatography (IEX-HPLC), and a novel two-dimensional high-performance liquid chromatography (2D-HPLC) method were used to detect the purity of bovine milk-derived exosomes with different processes of exosome isolation in detail. The results showed that the 2D-HPLC method could enhance the accuracy of detecting exosomal purity with higher precision and accuracy of instrumental, reduced personal error and experimental cost, shortened analysis time greatly, and more automation. The 2D-HPLC method is rapid, exhibits high selectivity, and has good sensitivity, thus making it well-suited for application in the pharmaceutical and toxicological analysis of exosomes.

## Introduction

Exosomes were considered garbage bags, carrying worthless cellular biomaterial [1–3]. At present, they are playing a crucial role in inter-cellular communication [4–8]. Nevertheless, the wide application of exosomes in drug delivery systems needed a better optimized isolation method for exosomes purification and cheaper source material [9]. In consideration of stability, nontoxicity, biocompatibility, tumor-targeting ability, and low cost, bovine milk-derived exosomes could become a kind of alternative natural drug delivery carrier [10–12].

Isolation method of exosomes that can give a higher yield with better purity and recovery of the exosomes play a crucial role in the collection of high◻quality exosomes for scaling up the operations in the industry [13–15]. Current isolation and characterization methods leave much to be desired in terms of ensuring high-quality exosomes. The most commonly used technologies for exosomes separation are based on physical properties, such as size and density, including ultracentrifugation (UC) [16, 17], density gradient centrifugation (DC) [18, 19], size exclusion chromatography (SEC) [20, 21], and ultrafiltration (UF) [22, 23]. The above methods, especially the use of DC achieved a much greater separation efficiency than the conventional method, thus providing milk exosomes of greater purity [24, 25], are closely followed by researchers. MISEV2018 noted that the degree of specificity of a particular method might vary depending on the type of biofluid from which EVs are separated. Hence, in our present study, we compared the differential physical and molecular characteristics of exosomes isolated using four different isolation protocols namely, the isolation by UC, DC, SEC, and UF. However, the current lack of quantitative methods of exosome purity, impedes our understanding of the true application values of exosomes.

Highly pure exosomes were essential for apply in various disease treatment fields. It was complicated to clearly define the content of each component because exosomes comprised various component types [26]. The typical characterization approaches of exosome purity included ratios of the different quantification methods may provide useful measures of purity. For example, protein:particle ratio [27, 28], protein:lipid ratio [29–31] and RNA:particle [32] have been proposed as possible purity metrics. At present, the main assessment methods of exosome purity were calculating the total particle number of exosomes per total protein (p/μg) or detecting the number of proteins of a particular size [33, 34]. The above conventions for measuring exosome purity were based on the quantity or size of proteins, so there were limitations to evaluating exosome purity. The measures for analyzing the total exosome particle amount were based on the Brownian motion of each particle by size [26, 35]. Absolute exosomes sizing and counting methods are currently imperfect and will require further improvement, aided by appropriate exosomes reference standards that are now in development [36]. In other words, in terms of quantification, precision and accuracy for exosome purity, the ratio between particle and protein are currently imperfect, and not the optimal method, which depends on exosome size [26]. Meanwhile, conventional detection methods of exosome purity remained disadvantageous, due to their high consumption of lab supplies and reagents, intricate equipment and experimental operation, high probability of artificial error, long detection time, and low accuracy. Therefore, according to the above-mentioned problems, we endeavored to establish an ideal method for characterization of exosome purity [13].

In the present study, we proposed an innovative a novel 2D-HPLC method for exosomal purity analysis by four different purification methods which reported that the purity of exosomes could be affected by different purification methods, combining size exclusion chromatography with ion exchange chromatography, that is a simple SE-HPLC analysis to detect watersoluble small-size proteins in samples, and an IEX-HPLC analysis to detect samples via similar size with different charges. 2D-HPLC method could potentially contribute significantly to studies on exosomes would lead to significant advances in the exosomes-based molecular diagnostics, treatment monitoring, drug delivery systems, test for drug resistance, and guide selection of therapeutic strategies.

## Materials and Methods

### Materials

Raw bovine milk (50-L/batch) was purchased from a local dairy factory (Zhongfen Dairying, Tianjin, China). Phosphate buffered saline (PBS) (10 ×) was purchased from Hyclone Laboratories. Inc. (USA) and diluted with ultrapure water (18.2 M Ω cm) obtained from a Millipore water system (USA). Analytical grade hydrochloric acid (HCl) and sodium hydroxide (NaOH)were purchased from Kermel (China). HPLC-grade trizma base (Tris) and Sodium chloride (NaCl) were purchased from Sigma. The above chemicals were used for experiments without further purification.

### Casein removal

HCL was added to the milk to adjust the pH to 4.6. The acidified milk stayed at room temperature for one hour to precipitate casein. The supernatant was collected after centrifugation at 4 °C and 4000 × g (Beckman Coulter, USA) for 30 min. The residual casein pellet was then removed by passing it through a 5-0.5 μm filtration system to obtain casein-free supernatant.

### Ultracentrifugation

Ultracentrifugation (UC) [16, 17] was a commonly used method for isolating exosomes. Nevertheless, it needed to consume several times and repetitive operations. The above casein-free supernatant was centrifuged to remove residual cellular components (4 °C, 100,000 × g, 105 min). The supernatant was transferred and repeated centrifugation three times by using the same method. Finally, the UC samples were stored at − 80°C before use.

### Density-gradient ultracentrifugation

Density-gradient ultracentrifugation [18, 19] was also a time-consuming (> 16 h) exosome separation technology. The casein-free supernatant was centrifuged at 4 °C, and 100,000 × g for 105 min to precipitate the bovine milk-derived EVs, which were subsequently resuspended in 1 mL PBS. The concentrated bovine milk-derived EVs were subjected to the top of a discontinuous density gradient consisting of 2 M, 1.65 M, 1.3 M, 0.95 M, and 0.6 M sucrose (7 mL volume for each gradient) in 250 mM Tris-HCl solution (pH 7.4). They were centrifuged at 100,000 × g at 4°C for 20 h. The bovine milk-derived exosome fractions between fraction 3 (1.3 M) and fraction 4 (1.65 M) were collected. To remove sucrose, the fraction was diluted in PBS to a final volume of 40 mL and centrifuged at 100,000 × g at 4 °C for 105 min. The pellet was resuspended in 1 mL PBS. Finally, the DC samples were stored at - 80 °C before use.

### Ultrafiltration

Ultrafiltration (UF) [22, 231] was a membrane separation technology. This purification method was based on the size and molecular weight of exosomes. The casein-free supernatant was tenfold condensed by centrifugation through 100 kDa molecular weight cut-off (MWCO) hollow fibers. Then the concentrate was washed in the same volume of PBS and centrifuged at 100,000 × g at 4 °C for 105 min. This washing step was repeated three times. Finally, the UF samples were stored at - 80°C before use.

### Size exclusion chromatography after ultrafiltration

Size exclusion chromatography (SEC) [20, 21] relied on the relative relationship between exosomal size and gel pore size. After the ultrafiltration process, the sample was purified using size exclusion chromatography (4FF chromatography packing, Bestarose, China). Wash the column with a double column volume of 0.5 M NaOH solution at 30 cm/h, followed by a double-column volume of water. The column was equilibrated with five times its volume using equilibration buffer (PBS, pH 8.0), and the sample loading volume was 5% of the column volume. The further purification sample was collected based on the determination of protein content by UV absorbance at 280 nm (> 50 mAU). Finally, the SEC samples were stored at −80°C before use.

### The particle size and number and total protein quantification

The particle size and number of bovine milk-derived exosome samples were characterized by the NanoFCM instrument (NanoFCM Inc., Xiamen, China) following the operations manual. A silica nanosphere cocktail (Cat. S16M-Exo, NanoFCM Inc., Xiamen, China) containing a mixture of 68 nm, 91 nm, 113 nm, and 155 nm standard beads was used to adjust the instrument for particle size measurement. The instrumental parameters were set as follows: Laser, 10 mW, 488 nm; SS decay, 10%; sampling pressure, 1.0 kPa; sampling period, 100 μs; Time to record, 1 min. The total protein content of exosomes was detected by the Pierce BCA Protein Assay Kit (Thermo Scientific, USA).

### Electron microscopy

The exosome morphology was studied by transmission electron microscopy (TEM). First, the bovine milk-derived exosomes sample (100 μg/mL) was fixed by mixing with an equal volume of 4% (w/v) paraformaldehyde at room temperature for 15 minutes. The fixed sample (10 μL) was then subjected to a formvar-carbon-coated TEM grid and kept at room temperature for 3 minutes. The grid was stained by adding 10 μL of uranyl oxalate solution (3% uranyl acetate, 0.0075 M oxalic acids, pH 7.0). The stained grid was investigated by a transmission electron microscope (Hitachi HT7800, Japan) operating at 120 KV.

### Western blotting

The detection of exosomal biomarkers was performed using a standard western blot procedure. The blots were hybridized with the following antibodies: TSG101 (BD Transduction Laboratories, USA), CD9 (Bio-Rad, USA), CD81, and Calnexin (Abcam, UK). Fluorescently conjugated rabbit or mouse antibodies were used as secondary antibodies. The films were visualized using the FluorChem E system (ProteinSimple, USA).

### Zeta potential

The electrokinetic potential of exosome samples was analyzed to assess particle stability in suspension. Exosomes samples were diluted certain ratio with PBS, and 500 μL samples diluted were loaded into disposable capillary cells, Brook zeta potential analyzer. The value was read in triplicates, and the average spectra were considered for comparative purposes. The low range was 0-28 V/cm, 1 to 250 Hz, sine or square wave, and the high range was 29-555 V/cm, same timing characteristics.

### Proteomics and bioinformatics analysis

The proteomics of bovine milk-derived exosomes (milk-exos) was analyzed by LC–MS/MS using Easy NLC 1200-Q Exactive Orbitrap mass spectrometers (ThermoFisher). The nano-HPLC system was equipped with an Acclaim PepMap nano-trap column (C18, 100 Å, 75 μm × 2 cm) and an Acclaim Pepmap RSLC analytical column (C18, 100 Å, 75 μm × 25 cm). Typically, 1 μL of the peptide mix was loaded onto the enrichment (trap) column at an isocratic flow of 5 μL/min of 2% CH3CN containing 0.1% formic acid for 5 min before the enrichment column was switched in-line with the analytical column. The eluents used for the LC were 2% CH3CN/0.1% (v/v) formic acid (solvent A) and 100% CH3CN/0.1% (v/v) formic acid (solvent B). The gradient used was 3% B to 25% B for 23 min, 25% B to 40% B in 2 min, 40% B to 85% B in 2 min, and maintained at 85% B for 2 min before equilibration for 10 min at 3% B before the next injection. All spectra were collected in positive mode using full-scan MS spectra scanning in the FT mode from m/z 300-1650 at resolutions of 70 000. A lock mass of 445.12003 m/z was used for both instruments. For MSMS on the QE, the 15 most intense peptide ions with charge states ≥2 were isolated with an isolation window of 1.6 m/z and fragmented by HCD with a normalized collision energy of 28. A dynamic exclusion of 30 seconds was applied.

The raw files were searched using Proteome Discover (version 2.1, ThermoFisher, Germany) with Sequest as the search engine. Fragment and peptide mass tolerances were set at 20 mDa and 10 ppm, respectively, allowing a maximum of 2 missed cleavage sites. The false discovery rates of proteins, peptides were 1 percent.

### Milk protein, IgM and IgG with ELISA determination

Quantitative detection of milk protein in bovine milk-derived exosomes was performed according to the manufacturer’s instructions (RIDASCREEN FAST Milk kit, R-Biopharm, Germany). Quantitative detection of IgM/IgG in exosomes was performed according to the manufacturer’s instructions (IgM/IgG cow ELISA kit, Abcam, UK). The absorbance was determined on a plate reader using the contents of each well at 450 nm. The average absorbance was calculated for standards and samples. The concentration of circulating milk protein, IgM and IgG was determined according to the standard curve obtained by plotting the mean absorbance for each standard concentration. Each sample was analyzed in duplicate using ELISA tests [37].

### HPLC with Size exclusion chromatography

The sample purity was analyzed by a column (4.6×150 mm, 2.5 μm, 450 Ǻ, Waters, USA) using a H-Class Ultra Performance Liquid Chromatography system (Waters, USA). The column was eluted at a flow rate of 0.5mL/min mobile phase (20 mM Tris-HCl, pH 7.2) with 25 μL injection of each sample. UV absorbance was detected at a wavelength of 280 nm.

### HPLC with anion exchange chromatography

The sample purity was analyzed by a column (1.3 μm, CIMac, DEAE 0.1 ml, BIA, Slovenia) using an H-Class Ultra-Performance Liquid Chromatography system (Waters, USA). The column was eluted at a flow rate of 0.6 mL/min mobile phase (phase A: 20 mM Tris-HCl, pH 7.2 and phase B: 20 mM Tris-HCl, 1 M NaCl, pH 7.2) with 25 μL injection of each sample. UV absorbance was detected at a wavelength of 280 nm.

### 2D-HPLC

The sample purity was first analyzed by a column (4.6×150 mm, 2.5 μm, 450 Ǻ, Waters, USA) using an H-Class Ultra-Performance Liquid Chromatography system (Waters, USA). The column was eluted at a flow rate of 0.5 mL/min mobile phase (20 mMTris-HCl, pH 7.2) with a 25 μL injection of each sample. UV absorbance was detected at a wavelength of 280 nm. And then the exosome fraction was injected into another column (1.3 um, CIMac, DEAE 0.1 ml, BIA, Slovenia) at a flow rate of 0.6 mL/min mobile phase (phase A: 20 mM Tris-HCl, pH 7.2, and phase B: 20 mM Tris-HCl, 1 M NaCl, pH 7.2). UV absorbance was detected at a wavelength of 280 nm.

## Results

### Preparation and characterization of bovine milk-derived exosomes by different purification technologies

Milk-exos were isolated and purified with DC, SEC, UF, and UC processes (Figure 1(a)). To validate whether the isolated extracellular vesicles (EVs) were exosomes, we observed the morphology by TEM. All of the corresponding TEM images showed the typical “cup-shaped” morphology of exosomes in all preparations (Figure 1(b)). For the four process, most of the size was 40-150 nm. The particle size of exosomes was assessed by nanoflow cytometry (Figure 1(c)). The mean particle size (the median particle size) of different processes’ exosomes was 73.39 nm (70.25 nm), 75.30 nm (71.75 nm), 74.02 nm (70.75 nm), and 72.95 nm (69.75 nm) for DC, SEC, UF, and UC, respectively. Moreover, the western blotting analysis shown that vital exosomal membrane markers CD9, CD81, and TSG101 were positive, and the microvesicle surface marker Calnexin was negative (Figure 1(d)), which confirmed that the isolated milk-exos were not contaminated with other multivesicular bodies. Meanwhile, the zeta potential value of exosomes shown in Figure 1(e) was −13.76 mV, −16.53 mV, −15.98 mV, and −15.08 mV for DC, SEC, UF, and UC, respectively.

**Figure 1.**
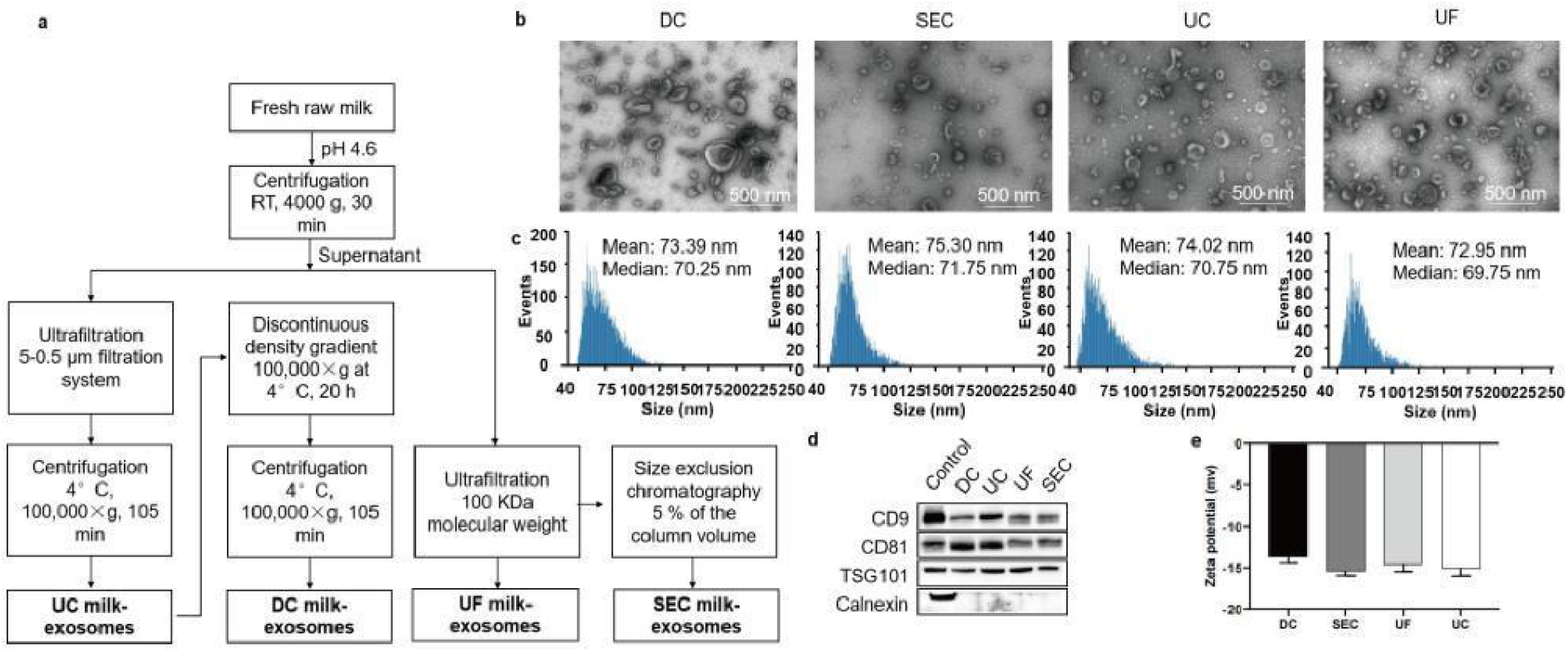
Characterisation of exosomes with DC, SEC, UF, UC processes. (a) Schematic representation of the major steps involved in isolating of exosomes from bovine raw milk with different processes. (b) The morphology and (c) size distribution profiles of exosomes particles with different processes was detected by TEM and nanoFCM, respectively. Scale bars = 500 nm. (d) Specific surface markers of exosomes were examined by western blot. The exosomes associated with markers were positive for CD9, CD81, and TSG101 and were negative for Calnexin. (e) Zeta potential value of exosomes with different processes were executed. Abbreviations: DC, density gradient ultracentrifugation; SEC, size exclusion chromatography after ultrafiltration; UF, ultrafiltration; UC, ultracentrifugation; TEM, transmission electron microscope.

To further analyze the protein components of exosomes, we carried out a proteomic analysis. As shown in Figure 2(a), according to the protein database search, the Venn diagram showed the overlap of proteins between different processes and most of the proteins; a total of 1730 proteins were identified in the exosomes with four different isolation methods. Exosomes from DC, SEC, UF, and UC processes had 1548, 1407, 1375, and 1288 proteins, respectively. Correlation analysis shown that in comparison with DC milk-exos, UC milk-exos, UF milk-exos, and SEC milk-exos protein quantities were significantly correlated with a Pearson r of 0.88, 0.86, and 0.92, respectively (Figure 2(b-d)). Proteomics analysis revealed the protein expression had different degrees of overlap between the four purification processes, and what’s more, the exosomes from other processes had significant differences in protein expression. 4.91% (76/1548), 7.11% (110/1548), and 4.26% (66/1548) protein expression for SEC, UF, and UC technologies of exosomes isolation was not coincidental with DC processes. According to the Venn diagram, different isolation methods influenced exosome quality.

**Figure 2.**
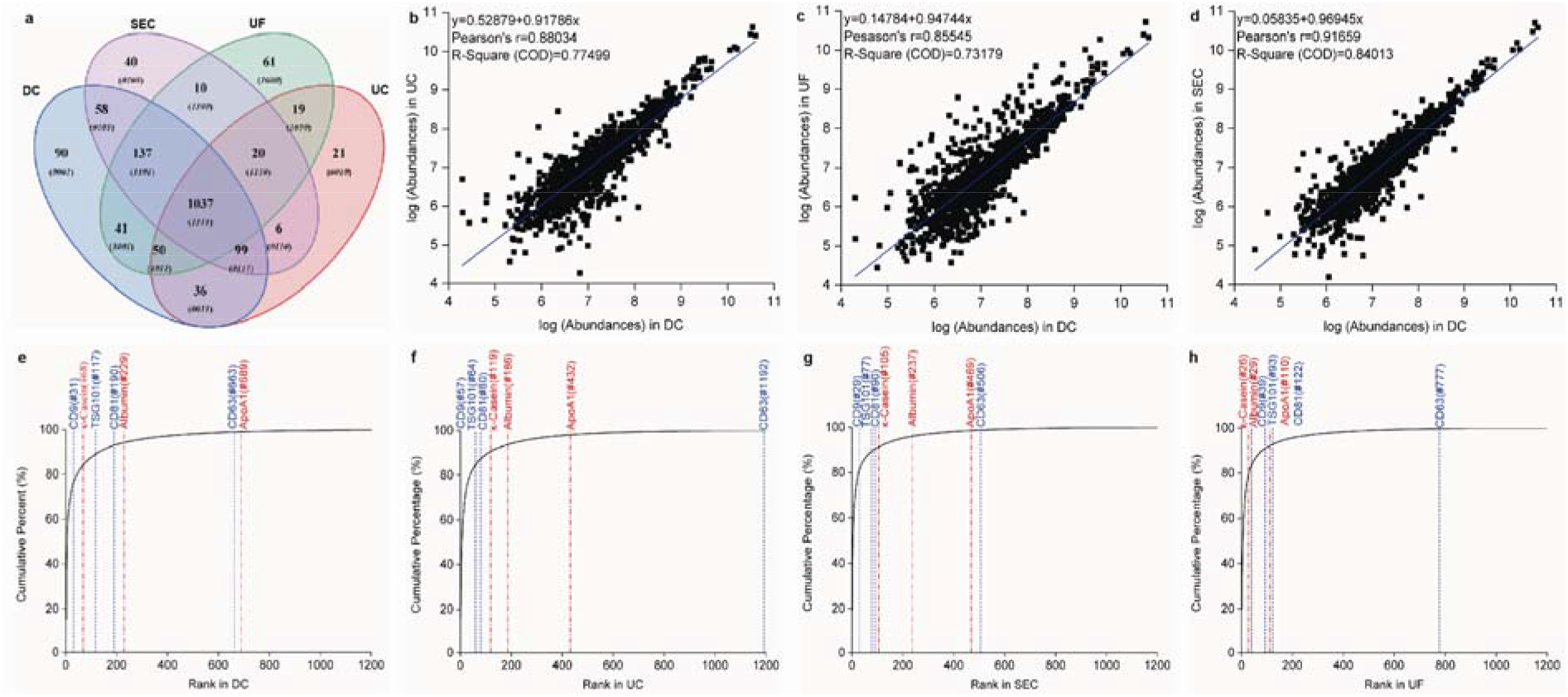
The comparison of exosomes with DC, SEC, UF, UC processes. (a) Venn diagram depicting the proteins expression between different processes. (b)-(d) Correlation analysis between different processes by Origin. (e)-(h) Protein quantity accumulation analysis between specific surface proteins of exosomes and major milk proteins (impurity protein). Specific surface proteins of exosomes include CD9, CD63, CD81, and TSG101 (blue), major milk proteins include κ-Casein, Albumin, and ApoA1 (red).

We then plotted cumulative curves of exosome protein quantities with DC, SEC, UF, and UC processes, and the quantities of proteins were ranked from high to low (Figure 2(e-h)). Exosome specific surface proteins including CD9, TSG101, CD81, and CD63, and major milk proteins including κ-Casein, Albumin, and ApoA1 were selected. Protein quantity accumulation analysis revealed exosome markers including CD81, CD9, CD63, and TSG101 that shared a similar ranking with DC, SEC, UF, and UC processes. While more contaminations were observed between UC milk-exos (κ-Casein, Albumin, and ApoA1 rank #119, #186, and #432) and UF milk-exos (κ-Casein, Albumin, and ApoA1 rank #26, #29, and #110). The major co-isolated contaminants were the bottom between DC (κ-Casein, Albumin, and ApoA1 rank #68, #229, and #689) and SEC processes (κ-Casein, Albumin, and ApoA1 rank #105, rank #237, rank #506).

### Analysis of exosome impurities and purities

The ratio of particle number to protein concentration was the most popular method used to evaluate the purity of exosomes. In this study, the protein concentration of exosomes was assessed using the BCA method. For four purification processes, the ratio of particle number to protein concentration was 1.40×10^11^, 0.30×10^11^, 0.14×10^11^, and 1.09×10^11^ particles per microgram for DC, SEC, UF, and UC, respectively (Table 1). The results suggested DC, SEC, and UC produced highly pure exosomes, exceeding the ratio of 3×10^10^, which was reported that the value of particle number to protein concentration over 3×10^10^ particles per microgram of protein was regarded as high purity, where the order of the particle-to-protein ratio was UC > DC > UF> SEC. However, the particle-to-protein ratio of UC was close to that of DC and far higher than SEC and UF, which was not in accordance with expectations. The reason for the abnormal result could be attributable to the limitation of this method depending on the quantity or size of the protein. Therefore, the purity method of the particle-to-protein ratio might be inaccurate to a certain extent.

**Table 1.** Analysis of exosome purities for the ratio of particle number to protein concentration.

|  | Particles<br>(10<sup>11</sup> Particles/ml) | Total protein<br>concentration<br>(μg/ml) | Ratio of particle<br>number to protein<br>concentration<br>(10<sup>11</sup> Particles/μg) |
| --- | --- | --- | --- |
| DC | 17.00 | 12.15 | 1.40 |
| SEC | 4.32 | 14.43 | 0.30 |
| UF | 7.81 | 56.92 | 0.14 |
| UC | 44.20 | 40.61 | 1.09 |

Beyond that, ELISA was a method for evaluating exosome purity by detecting impurity concentration. Hence, we evaluated exosome purity with ELISA technology, including milk protein, IgG and IgM, which were shown in Table 2. For the content of three impurities, both DC and SEC were lower than UF or UC. Furthermore, whatever the impurity was, the impurity content of DC far less than the other processes. The difference in impurity concentration between UF and UC was relatively small, where the order of impurity concentration was UF > UC > SEC> DC.

**Table 2.**
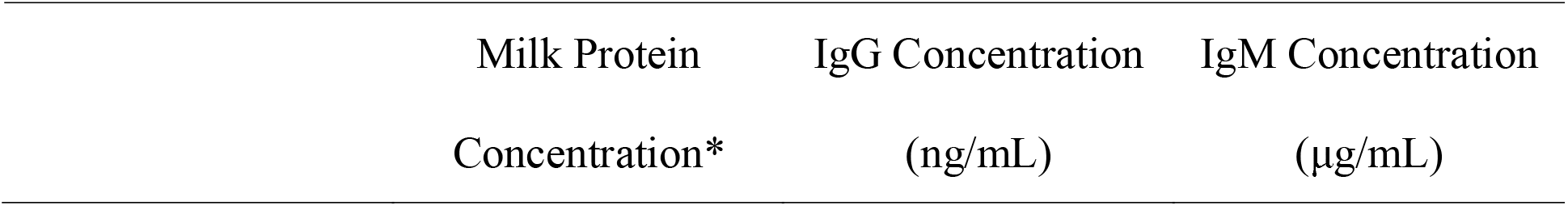
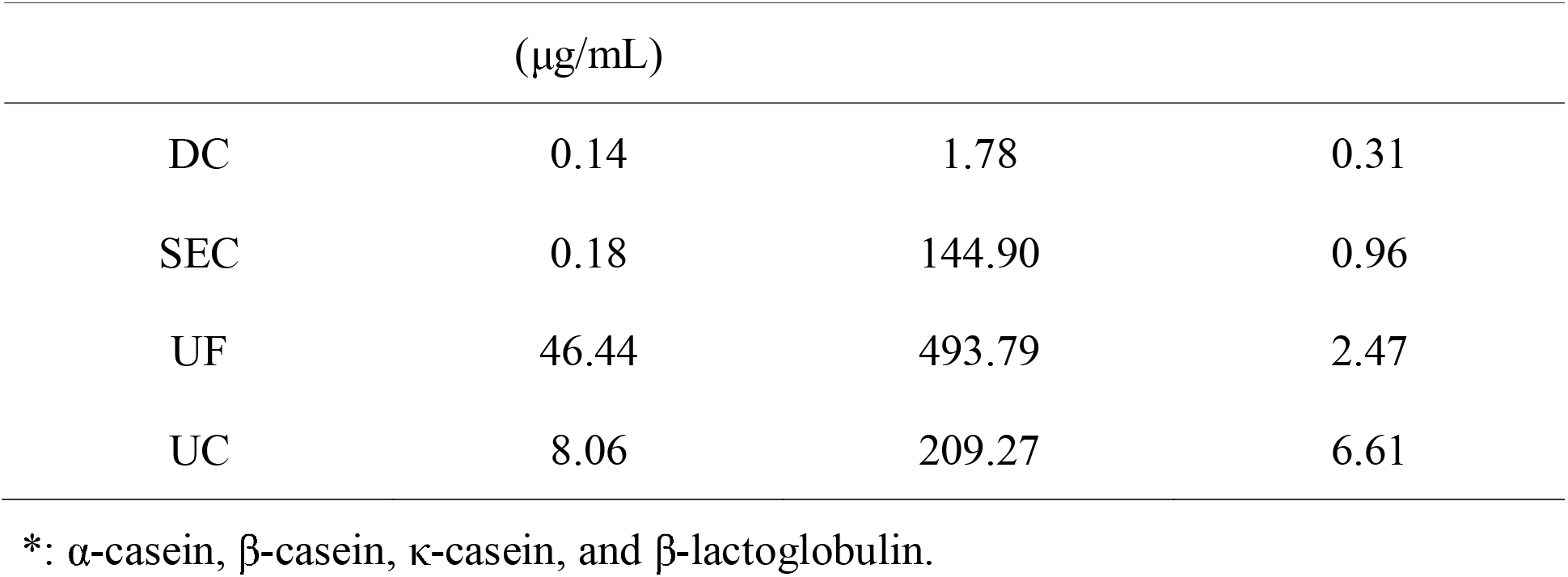
Analysis of exosome impurities for the ELISA assay.

### Purity analysis of bovine milk-derived exosomes by the SE-HPLC method

Selection of the most appropriate prepacked column or packing material is crucial for success, since the support pore size controls the solute size range in which fractionation occurs. SEC has better separation (based on size) and helps to eliminate contaminants with more confidence. As the suitable SEC columns for packing, 450 Å column, 1000 Å column, and 2000 Å column were selected to analysis the exosome putity according to pore size. The purity analysis of exosomes was assessed by size-exclusion high-performance liquid chromatography (SE-HPLC) with 450 Å column, 1000 Å column, and 2000 Å column, respectively (Figure 3). The purity of exosomes purified by DC, SEC, UF, and UC was 99.29%, 94.11%, 78.05%, and 79.29%, respectively, with a 450 Å column (Figure 3(a-d)), while the purity of exosomes purified by DC, SEC, UF, and UC both was 100% with a 1000 Å column (Figure 3(e-h)) and a 2000 Å column (Figure 3(i-l)). As illustrated in Figure 3(a-d), the relative retention time of the principal peak was about 1.6 min, whichever the isolation process of exosomes was, suggesting the favorable stability of the SE-HPLC analysis method. The HPLC chromatogram of different technological exosomes appeared with 1-3 impurity peaks. Furthermore, the purity of exosomes with DC and SEC was considerably higher than the other processes, which demonstrated exosomes isolated by UF and UC remained contaminated with abundant impurities. Exosomes with SEC had less impurity, and DC possessed the best purity of the four processes.

**Figure 3.**
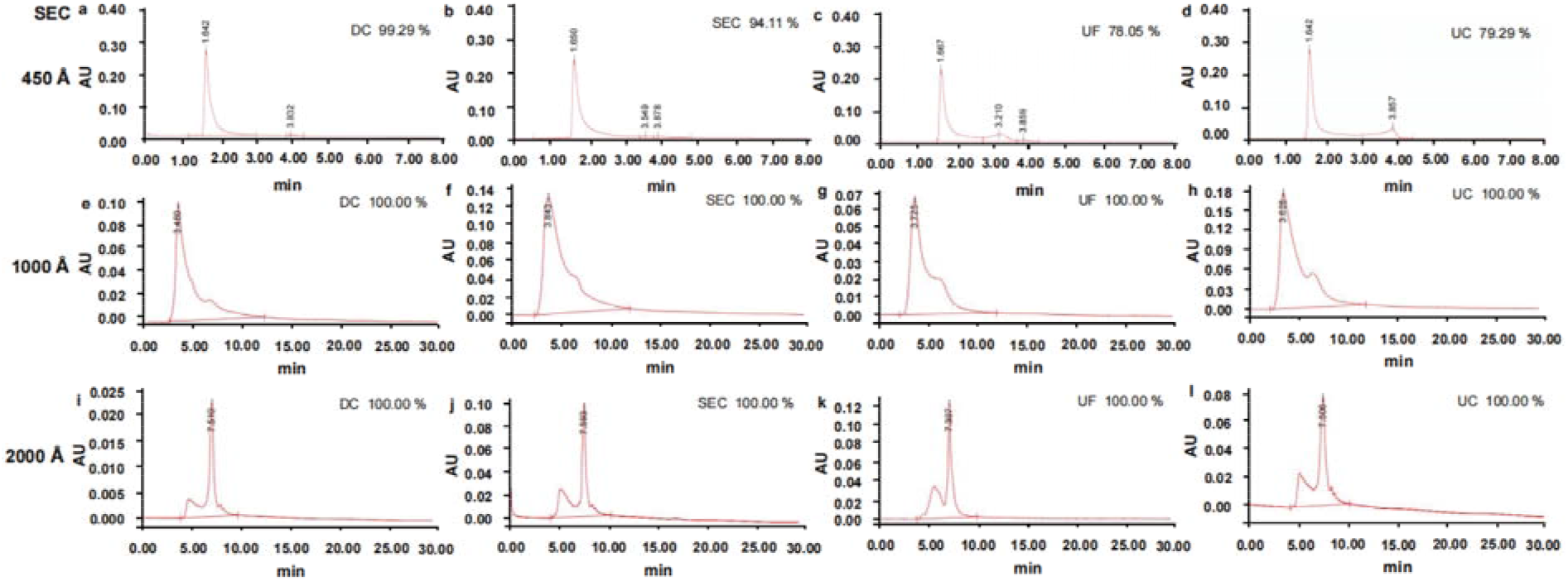
Purity analysis of exosomes by the SE-HPLC method. Exosomal chromatograms of (a), (e), and (i) DC, (b), (f), and (j) SEC, (c), (g), and (k) UF, and (d), (h), and (l) UC were analyzed with 450 Å column, 1000 Å column, 2000 Å column, respectively.

### Purity analysis of bovine milk-derived exosomes by the IEX-HPLC method

Proteins with highly similar sizes cannot be separated by SEC, but if they have diferent pI values they can be separated by IEX chromatography. Though the SE-HPLC analysis method could detect many impurities according to different size, it was difficult to distinguish exosomes from other impurities of similar size. Therefore, the chromatogram of ion-exchange high-performance liquid chromatography (IEX-HPLC) was used to analyze the purity of exosomes becsuse of charge heterogeneity (Patent NO. CN113533589A). Figure 4 revealed the chromatogram of ion-exchange high-performance liquid chromatography (IEX-HPLC) of exosomes with different purification technologies. As illustrated in Figure 4, proteins bind the stationary phase based on their overall charge, and the density and distribution of their surface charge. Elution of the bound proteins from the column is performed via salt gradient, such as NaCl. A weakly bound protein elutes from the matrix at low salt concentrations (low conductivity) and strongly bound protein elutes at higher salt concentrations (high conductivity). Charge heterogeneity results discovered the peaks of charge-heterogeneous exosomes by IEX-HPLC could be separated by optimizing the gradient towing conditions (Figure 4(a-d)). As shown in Figure 4(e), the chromatogram of DC arose six peaks due to heterogeneity of charge. And the six peaks appeared in each exosome chromatogram of different processes. According to the result of the purity analysis by SE-HPLC, DC has the highest purity and hardly any impurities. Therefore, the six peaks were considered as the main peaks, and the exosome purity of DC was regarded as 100.00%, making it a “reference substance”. The chromatogram of the other processes showed a new peak around 4.3 min, which was the impurity peak. The purity of SEC, UF, and UC was 96.49%, 94.40%, and 93.03%, respectively (Figure 4(f-h)). The purity differences of exosomes were not obvious in the different processes. Furthermore, the purity of UF and UC was quite high, which might be inaccurate. Besides, characterization of the six peaks with the exosome purity from DC showed that the morphology of exosomes exhibited a rich profusion of exosomes consistent with classical exosome-like morphology, size distribution and protein markers (Supplementary Figure 1).

**Figure 4.**
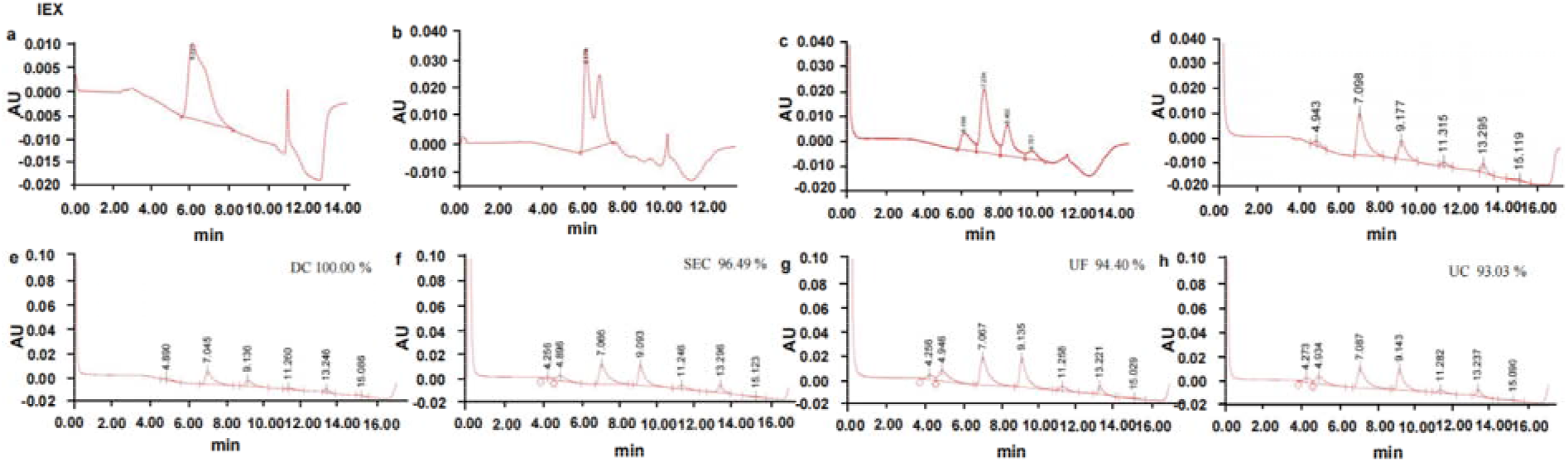
Purity analysis of exosomes by the IEX-HPLC method. (a)-(d) Charge heterogeneity discovery results of exosomes by IEX-HPLC. Exosomal chromatogram of (e) DC, (f) SEC, (g) UF, and (h) UC was detected.

### A novel purity analysis method of bovine milk-derived exosomes by 2D-HPLC

The IEX-HPLC method could analyze exosomal impurities based on charge heterogeneity, but only small impurities could be detected. To take full advantage of SE-HPLC and IEX-HPLC, we assessed exosome purity by using two-dimensional high-performance liquid chromatography (2D-HPLC), which was the SE-HPLC method combined with the IEX-HPLC method. Figure 5 shown the exosome chromatogram of 2D-HPLC, that primary-dimensional HPLC adopted the SE-HPLC method and second-dimensional HPLC adopted the IEX-HPLC method. The final purity was obtained by multiplying the purity in primary-dimensional HPLC and second-dimensional HPLC together. Figure 5(a) suggested the exosome purity of DC (100.00%) was pure enough whether in primary-dimensional or second-dimensional HPLC. For SEC process, the final purity was 76.15% which was the product of multiplying the purity in primary-dimensional HPLC (77.75%) or second-dimensional HPLC (97.94%). For the UF process, the final purity was 67.47% which was the product of multiplying the purity in primary dimensional (71.64%) or second-dimensional HPLC (94.18%). For the UC process, the final purity was 65.96% which was the product of multiplying the purity in primary dimensional (71.40%) or second-dimensional HPLC (92.39%) (Figure 5). The chromatogram of 2D-HPLC was in agreement with the chromatography of SE-HPLC (Figure 3) and IEX-HPLC (Figure 4) separately, suggesting the 2D-HPLC method was successfully compatible.

**Figure 5.**
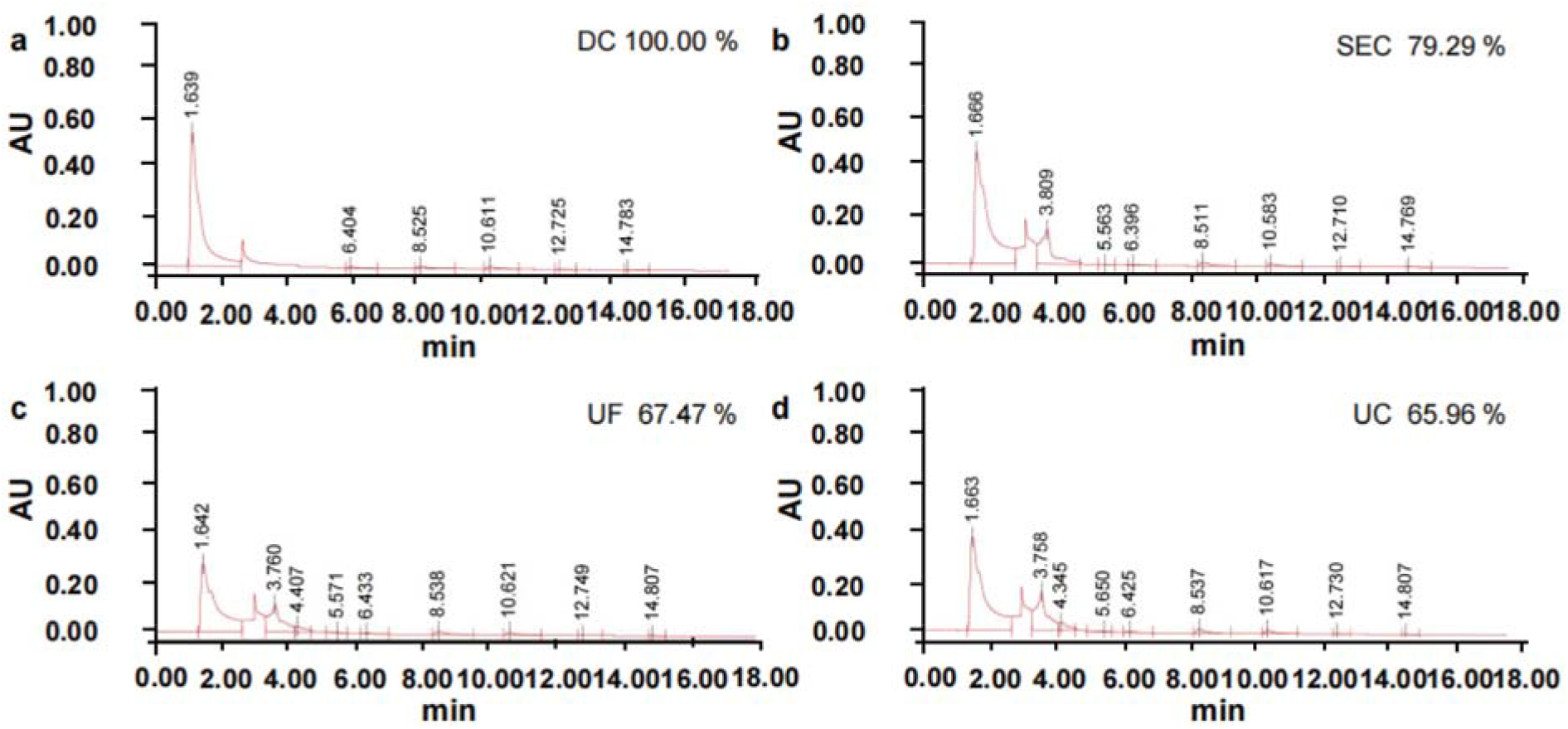
Purity analysis of exosomes by the 2D-HPLC method. Exosomal chromatogram of (a) DC, (b) SEC, (c) UF, and (d) UC was executed.

## Discussion

In this study, we first successfully developed a qualitative and quantitative method, a novel two-dimensional high-performance liquid chromatography, to analyze the purity of milk exosome isolated from different purification technologies. Such studies contribute to our understanding of the application of advanced analytical technologies.

So far, most research relied on several technologies to evaluate exosomal purity comprehensively. However, it might still be inaccurate. For example, the ratio of particle number to protein concentration was the most popular method to assess the purity of exosomes. Nonetheless, the method had certain limitations. The detection of particle number relied on the Brownian motion of each particle by size [26, 35]. The accuracy of experimental equipment was easily influenced by some parameters and operation conditions. Furthermore, the method was difficult to use for detecting small impurities [15, 38, 39]. As a result, this method was unsuitable for use as a quality control criterion method. We urgently needed a purity method to assess the quality of exosomes in large-scale manufacturing. Other analysis technologies for exosomal purity, such as ELISA, WB, qPCR, and so on, also have some limitations. These means had a high consumption of lab supplies and reagents, intricate equipment and experimental operations, a high probability of artificial error, a long detection time, and low accuracy. These limitations were unfavorable for detecting the purity of exosomes quickly, and waiting for the purity result would delay the release of the finished product. It was no doubt that the wait time increased production costs. For this reason, it was crucial to find a suitable method of assessing exosomal purity.

To take full advantage of SE-HPLC and IEX-HPLC methods, we tried to develop a novel analysis method of 2D-HPLC combining both two methods. Primary-dimensional HPLC uses the SE-HPLC method, and second-dimensional HPLC adopted the IEX-HPLC method. The final purity was the result of multiplying the purity in primary-dimensional HPLC and second-dimensional HPLC together. The fraction of principal peak only, flowing through primary dimensional HPLC by using SE-HPLC method, entered into second-dimensional HPLC adopted IEX-HPLC method which performed a secondary analysis. The results showed that the SE-HPLC analysis method could detect many impurities according to sizes, while it was challenging to distinguish exosomes from other impurities with similar sizes; the IEX-HPLC method could analyze exosomal impurities based on charge heterogeneity, but only a few impurities could be detected; and the 2D-HPLC method, taking full advantage of the SE-HPLC and IEX-HPLC methods, could enhance the accuracy of detecting exosomal purity relative to SE-HPLC, improving the disadvantage of SE-HPLC to some extent. Our results also showed that 2D-HPLC had the amount of advantages relative to other analysis means of exosomal purity, such as higher precision and accuracy of instrumental, reduced personal error and experimental cost, shortened analysis time greatly, and more automation. These advantages contributed to the realization that the 2D-HPLC method became an exosomal analysis method for quality control criteria. It should be noted that the inability to characterize impurities, a professional operation, and intricate equipment are the limitations of 2D-HPLC. In this study, we researched the analysis methods for purity, which focused on bovine milk-derived exosomes. In subsequent investigations, we aim to utilize the assay established in the present study to examine and contrast exosomes obtained from diverse origins, thereby enhancing the resolution and accuracy of this analytical methodology. Additionally, this will facilitate a more comprehensive understanding of exosomes and offer a valuable reference for researchers working in the field of exosomes.

## Acknowledgements

We thank Mei Bai for helpful discussions.

## Author contributions

\# L.L., C.L.H., Q.Z. contributed equally.

## Data Availability Statement

All data are available in the main text or the supplementary materials. Further inquiries can be directed to the corresponding author.

## References

[1] Kanchanapally R, Deshmukh S K, Chavva S R, et al. Drug-loaded exosomal preparations from different cell types exhibit distinctive loading capability, yield, and antitumor efficacies: a comparative analysis. International journal of nanomedicine, 2019, 14: 531.

[2] Théry C, Zitvogel L, Amigorena S. Exosomes: composition, biogenesis and function. Nature reviews immunology, 2002, 2(8): 569–579.

[3] Kalluri R. The biology and function of exosomes in cancer. The Journal of clinical investigation, 2016, 126(4): 1208–1215.

[4] Février B, Raposo G. Exosomes: endosomal-derived vesicles shipping extracellular messages. Current opinion in cell biology, 2004, 16(4): 415–421.

[5] Thakur B K, Zhang H, Becker A, et al. Double-stranded DNA in exosomes: a novel biomarker in cancer detection. Cell research, 2014, 24(6): 766–769.

[6] Janas T, Janas M M, Sapoń K, et al. Mechanisms of RNA loading into exosomes. FEBS letters, 2015, 589(13): 1391–1398.

[7] Bang C, Thum T. Exosomes: new players in cell–cell communication. The international journal of biochemistry & cell biology, 2012, 44(11): 2060–2064.

[8] Whiteside T L. Exosomes carrying immunoinhibitory proteins and their role in cancer. Clinical & Experimental Immunology, 2017, 189(3): 259–267.

[9] Singh A, Sreenu B, Alvi S, et al. Bovine milk derived exosomal-curcumin exhibiting enhanced stability, solubility, and cellular bioavailability. Clinical Oncology, 2021, 6: 1769.

[10] Oskouie M N, Aghili Moghaddam N S, Butler A E, et al. Therapeutic use of curcumin◻encapsulated and curcumin◻primed exosomes. Journal of cellular physiology, 2019, 234(6): 8182–8191.

[11] Sun D, Zhuang X, Xiang X, et al. A novel nanoparticle drug delivery system: the anti-inflammatory activity of curcumin is enhanced when encapsulated in exosomes. Molecular therapy, 2010, 18(9): 1606–1614.

[12] Kim M S, Haney M J, Zhao Y, et al. Development of exosome-encapsulated paclitaxel to overcome MDR in cancer cells. Nanomedicine: Nanotechnology, Biology and Medicine, 2016, 12(3): 655–664.

[13] Contreras-Naranjo J C, Wu H J, Ugaz V M. Microfluidics for exosome isolation and analysis: enabling liquid biopsy for personalized medicine. Lab on a Chip, 2017, 17(21): 3558–3577.

[14] Shao H, Chung J, Lee K, et al. Chip-based analysis of exosomal mRNA mediating drug resistance in glioblastoma. Nature communications, 2015, 6(1): 1–9.

[15] Huang T, Banisz A B, Shi W, et al. Size exclusion HPLC detection of small-size impurities as a complementary means for quality analysis of extracellular vesicles. Journal of circulating biomarkers, 2015, 4(Godište 2015): 4–6.

[16] Kim H, Shin S. ExoCAS-2: rapid and pure isolation of exosomes by anionic exchange using magnetic beads. Biomedicines, 2021, 9(1): 28.

[17] Lin S, Yu Z, Chen D, et al. Progress in microfluidics◻based exosome separation and detection technologies for diagnostic applications. Small, 2020, 16(9): 1903916.

[18] Zhang M, Jin K, Gao L, et al. Methods and technologies for exosome isolation and characterization. Small Methods, 2018, 2(9): 1800021.

[19] Kamerkar S, LeBleu V S, Sugimoto H, et al. Exosomes facilitate therapeutic targeting of oncogenic KRAS in pancreatic cancer. Nature, 2017, 546(7659): 498–503.

[20] Batrakova E V, Kim M S. Using exosomes, naturally-equipped nanocarriers, for drug delivery. Journal of Controlled Release, 2015, 219: 396–405.

[21] Lane R E, Korbie D, Trau M, et al. Optimizing size exclusion chromatography for extracellular vesicle enrichment and proteomic analysis from clinically relevant samples. Proteomics, 2019, 19(8): 1800156.

[22] Quintana J F, Makepeace B L, Babayan S A, et al. Extracellular Onchocerca-derived small RNAs in host nodules and blood. Parasites & vectors, 2015, 8(1): 1–11.

[23] Alvarez M L, Khosroheidari M, Ravi R K, et al. Comparison of protein, microRNA, and mRNA yields using different methods of urinary exosome isolation for the discovery of kidney disease biomarkers. Kidney international, 2012, 82(9): 1024–1032.

[24] Thery C, Witwer KW, Aikawa E, et al. Extracell J. Vesicles 2018, 7, 1535750.

[25] Witwer KW, Buzas EI., Bemis LT, et al. Standardization of sample collection, isolation and analysis methods in extracellular vesicle research. J. Extracell. Vesicles, 2013, 2(1): 20360.

[26] Ahn S H, Ryu S W, Choi H, et al. Manufacturing Therapeutic Exosomes: from Bench to Industry. Molecules and Cells, 2022, 45(5): 284.

[27] Webber J, Clayton A How pure are your vesicles? J Extracell Vesicles, 2013, 2: 19861.

[28] Maiolo D, Paolini L, Di Noto G, et al. Colorimetric nanoplasmonic assay to determine purity and titrate extracellular vesicles. Anal Chem, 2015, 87(8): 4168–4176.

[29] Osteikoetxea X, Balogh A, Szabó-Taylor K, et al. Improved characterization of EV preparations based on protein to lipid ratio and lipid properties. PLoS One. 2015;10(3): e0121184.

[30] Mihály J, Deák R, Szigyártó IC, et al. Characterization of extracellular vesicles by IR spectroscopy: fast and simple classification based on amide and CH stretching vibrations. Biochim Biophys Acta, 2017, 1859(3): 459–466.

[31] Lai RC, Arslan F, Lee MM, et al. Exosome secreted by MSC reduces myocardial ischemia/reperfusion injury. Stem Cell Res, 2010, 4(3): 214–222.

[32] Cvjetkovic A, Lotvall J, Lasser C The influence of rotor type and centrifugation time on the yield and purity of extracellular vesicles. J Extracell Vesicles, 2014, 3: 23111.

[33] Webber J, Clayton A. How pure are your vesicles? Journal of extracellular vesicles, 2013, 2(1): 19861.

[34] Angelbello A J, Benhamou R I, Rzuczek S G, et al. A small molecule that binds an RNA repeat expansion stimulates its decay via the exosome complex. Cell chemical biology, 2021, 28(1): 34–45. e6.

[35] Bachurski D, Schuldner M, Nguyen P H, et al. Extracellular vesicle measurements with nanoparticle tracking analysis–An accuracy and repeatability comparison between NanoSight NS300 and ZetaView. Journal of extracellular vesicles, 2019, 8(1): 1596016.

[36] Valkonen S, van der Pol E, Böing A, et al. Biological reference materials for extracellular vesicle studies. Eur J Pharm Sci, 2017, 98: 4–16.

[37] Turlej E, Goszczyński T M, Drab M, et al. The Impact of Exosomes/Microvesicles Derived from Myeloid Dendritic Cells Cultured in the Presence of Calcitriol and Tacalcitol on Acute B-Cell Precursor Cell Lines with MLL Fusion Gene. Journal of clinical medicine, 2022, 11(8): 2224.

[38] Gardiner C, Ferreira Y J, Dragovic R A, et al. Extracellular vesicle sizing and enumeration by nanoparticle tracking analysis. Journal of extracellular vesicles, 2013, 2(1): 19671.

[39] Kang Y T, Kim Y J, Bu J, et al. High-purity capture and release of circulating exosomes using an exosome-specific dual-patterned immunofiltration (ExoDIF) device. Nanoscale, 2017, 9(36): 13495–13505.

