## Supplemental Figure 1 for "A novel purity analysis method of bovine milk-derived exosomes by two-dimensional high-performance liquid chromatography"

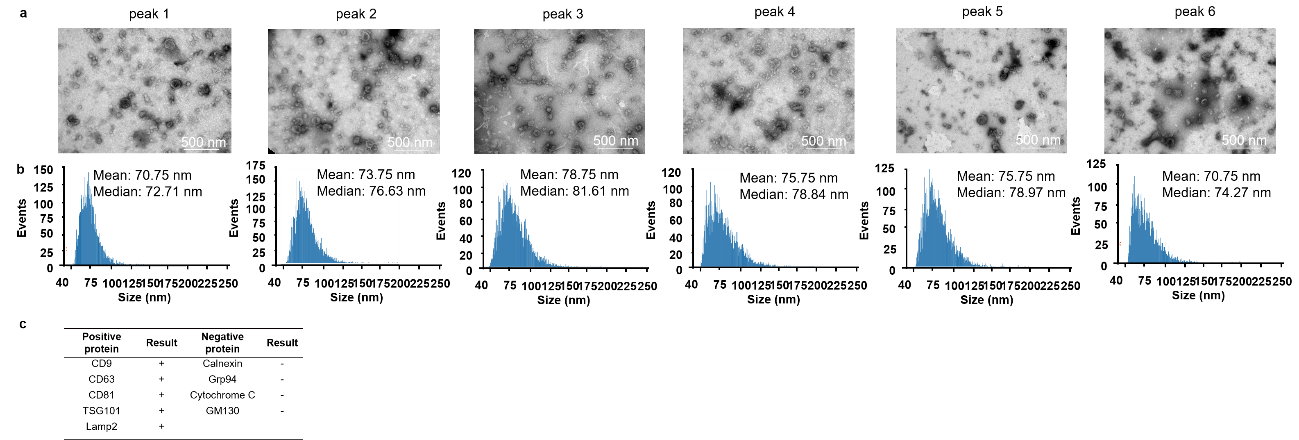

**Supplementary Figure 1**. **Characterisation of exosomes with different peak of SE-HPLC.** (a) The morphology and (b) size distribution profiles of exosomes particles with six peaks of SE-HPLC were detected by TEM and nanoFCM, respectively. Scale bars = 500 nm. (c) Specific surface markers of exosomes were examined by proteomic analysis. The exosomes associated with markers were positive for CD9, CD81, CD63, Lamp2, and TSG101 and were negative for Calnexin, GM130, Grp94, and Cytochrome C.
